# The polyploid state plays a tumor suppressive role in the liver

**DOI:** 10.1101/149799

**Authors:** Shuyuan Zhang, Keijin Zhou, Xin Luo, Lin Li, Liem Nguyen, Yu Zhang, Branden Tarlow, Daniel Siegwart, Hao Zhu

## Abstract

Most cells in the liver are polyploid, but the functional role of polyploidy is unknown. Polyploidization normally occurs through cytokinesis failure and endoreduplication around the time of weaning. To interrogate the function of polyploidy while avoiding irreversible manipulations of essential cell cycle genes, we developed multiple orthogonal mouse models to transiently and potently alter liver ploidy. Premature weaning, as well as in vivo knockdown of *E2f8* or *Anln*, allowed us to toggle between diploid and polyploid states. While there was no impact of ploidy alterations on liver function, metabolism, or regeneration, hyperpolyploid mice suppressed and hyperdiploid mice accelerated tumorigenesis in mutagen and high fat induced models. Mechanistically, the diploid state was more susceptible to Cas9-mediated tumor suppressor loss but was similarly susceptible to *MYC* oncogene activation, indicating that ploidy differentially protected the liver from distinct genomic aberrations. Our work suggests that polyploidy evolved to prevent malignant outcomes of liver injury.

## Introduction

Polyploid cells and organisms contain more than two homologous sets of chromosomes. Polyploidy is prevalent in plants, fish, and salamanders^1^, but rare in mammals except in the cells of the heart, marrow, and liver. Up to 90% of rodent and 50% of human hepatocytes are polyploid^2,3^. In rodents, liver ploidy dramatically increases around weaning (P14-P21) and continues to increase with age^4,5^. The dominant mechanism for polyploidization is cytokinesis failure that leads to binucleated hepatocytes^5^. To a lesser extent, hepatocyte endoreduplication also contributes to polyploidy via replication of the nuclear genome in the absence of cell division^6^. Thus, polyploid hepatocytes (tetraploids, octoploids, etc.) can be bi- or mononuclear. Postnatal liver polyploidization is developmentally regulated, but ploidy is dynamic and can increase with surgery^7^, fatty liver disease, and oxidative stress^8^. Despite these observed correlations, the extent to which ploidy influences cellular or tissue function remains completely unknown.

Many hypotheses have been proposed for the existence of polyploid cells. In yeast and plants, polyploidy promotes adaptation to environmental stresses, in some cases through the production of genetically diverse aneuploid daughter cells^9,10^. Recently, it was shown that a subset of polyploid hepatocytes undergo reductive cell divisions prone to missegregation, leading to the accumulation of aneuploid cells^11^. Though the prevalence of aneuploidy in the liver is debated^12^, aneuploidy may represent a means by which genetic diversity and premalignancy can evolve. Whether or not polyploidy is a risk for or protective against cancer is unknown. In support of the premalignancy concept, Fujiwara *et al.* showed that tetraploidy in *p53* null mouse mammary epithelial cells is a chromosomally unstable state that predisposes to transformation, but the relevance to wild-type tetraploid hepatocytes is unclear^13^. As an argument against increased cancer risk, the polyploid state has been associated with terminal differentiation and limited proliferation^14^. Also, Ganem *et al.* demonstrated that tetraploid hepatocyte proliferation/transformation is suppressed by the Hippo pathway^15^, but we and others have shown that 4c and 8c hepatocytes divide efficiently and contribute as much to proliferation, growth, and regeneration as 2c cells in vivo (Extended Data Fig. 1a)^11^. Despite the hypothetical risks and benefits of polyploidy, whether tetraploid or diploid hepatocytes are more prone to cancer remains untested and thus the functional contributions of polyploidy to liver physiology, injury, and cancer remain completely unknown.

Defects in genes required for cell cycle or cytokinesis (*Trp53, Rb*, or *Cdk1)* can dramatically alter ploidy^16^^-^^18^, but interrogation of mice with these germline mutations cannot distinguish the effects of ploidy from the effects of profound cell cycle deficiencies. For example, *Cdk1* knockout (KO) hepatocytes cannot complete mitosis and are dramatically polyploid (up to 32N or greater). As a consequence, liver-specific *Cdk1* KO mice are unable to undergo malignant transformation, but whether this is specifically due to polyploidy or permanent *Cdk1* deficiency is unclear^18^. In contrast, *E2f7*/*E2f8* double KO livers are almost entirely comprised of diploid hepatocytes, but these mice have no physiological or regenerative phenotypes^19,20^, suggesting that polyploidy is dispensable for liver development and regeneration after acute damage. Recently, *E2f8* KO livers were shown to accelerate tumorigenesis, but it is unknown if this is due to transcriptional effects of *E2f8* deficiency or to the diploid state of *E2f8* KO hepatocytes^21^. Likewise, Zhang et al. showed that *Yap* activation promotes polyploidy and cancer, but it is unclear if the ploidy itself directly impacts tumor biology independent of *Yap*^22^. Due to the lack of appropriate tools and the inability to assess the chronic impact of ploidy change, the role polyploidy plays in diseases involving long-term cell division cycles remains unclear. Here, we developed multiple methods and in vivo reagents to transiently, reversibly alter ploidy and found that the polyploid state suppressed liver tumorigenesis by buffering against tumor suppressor loss.

### Premature weaning promoted polyploidy and was protective against HCC development

To answer these questions, we reasoned that the transient, reversible control of ploidy state would represent a fundamental advance to enable the elucidation of ploidy functions. Since the predominant mechanism of widespread hepatocyte polyploidization is cytokinesis failure, a phenomenon temporally associated with weaning in rats^23^, we asked if differential weaning times could transiently influence ploidy in mice. By weaning wild-type (WT) mice at P13 (premature weaning) or P21 (normal weaning) (Fig. 1a), we found that prematurely weaned mice had significantly more binucleated hepatocytes (Fig. 1b) and increased cellular ploidy (Fig. 1c) at 19 days of age. To induce hepatocellular carcinoma (HCC), we gave a single intraperitoneal (IP) dose (25μg/g) of the agent diethylnitrosamine (DEN) to both cohorts at P19, a time point when ploidy states were divergent. Five months later, pre-weaned mice with greater ploidy at P19 exhibited significantly reduced gross and microscopic tumor burden (Fig. 1d, e), suggesting that polyploidy could exert tumor suppressive effects. Since weaning is confounded by factors other than ploidy, we also used additional methods to control ploidy state.

**Figure 1.**
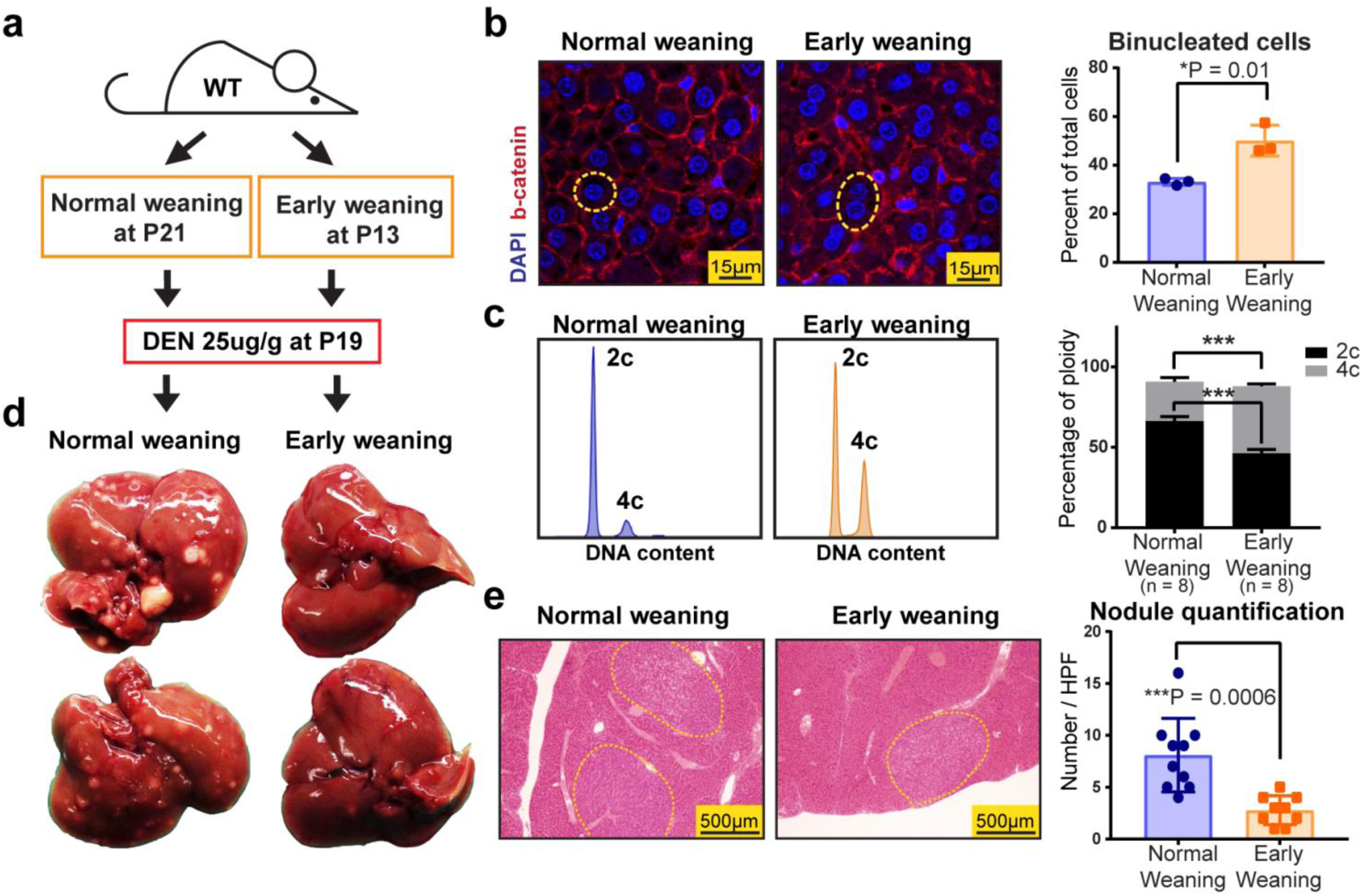
Premature weaning promoted polyploidy and was protective against HCC development. a. WT mice were weaned at P13 and P21 to induce differences in polyploidization. Diethylnitrosamine (DEN) was injected to both cohorts at P19 to induce hepatocellular carcinoma (HCC) formation.
b. DAPI and Ctnnb1 immunofluorescence staining identified nuclei and cell membranes (left panel). Representative mononucleated and binucleated hepatocytes are circled by the yellow dashed lines. The percentages of binucleated hepatocytes are quantified on the right (4 images were analyzed and averaged per mouse, n = 3 mice per group).
c. Ploidy distribution was analyzed by flow cytometry. Representative histograms show DNA content as stained by PI. Compiled data from the two groups are shown on the right (n = 8 mice in each group).
d. Gross tumor burden was examined five months after DEN.
e. Microscopic tumor nodules are circled by yellow dashed lines. The right panel shows nodule quantification (One section of an entire liver lobe was quantified per mouse, n = 9 mice per group).

### In vivo siRNAs revealed that the polyploid state protected against tumorigenesis

To inducibly toggle polyploidy without introducing permanent genetic lesions, we used siRNA to transiently knockdown genes that affect ploidy. We targeted *Anillin (Anln)*, an actin binding protein required for cytokinesis^24^ (Extended Data Fig. 1b, c) and *E2f8*, a transcription factor required for polyploidization^20^. In vivo, *Anln* and *E2f8* siRNA vs. *scramble* siRNA (siCtrl) delivery in lipid nanoparticles from P10-P20^25^ resulted in significant knockdown of *Anln* and *E2f8* mRNA and protein (Fig. 2a, b and Extended Data Fig. 1d). As expected, cellular ploidy was significantly increased after *Anln* knockdown and decreased after *E2f8* knockdown (Fig. 2c). Confocal imaging based ploidy characterization revealed that siAnln treated livers had significantly larger cell and nuclear size, while the siE2f8 treated livers showed the opposite (Fig. 2d, e). Since cell and nuclear size correlate with DNA content, this suggested significant ploidy changes. To quantitatively distinguish mono- vs. binucleated tetraploids, we integrated flow and confocal imaging data (Fig. 2f). This revealed that siAnln hepatocytes were 32% mononuclear diploid, 32% mononuclear tetraploid, and 32% binuclear tetraploid. siE2f8 hepatocytes were 64% mononuclear diploid, 17% mononuclear tetraploid, and 16% binuclear tetraploid. Furthermore, a Florescence In Situ Hybridization (FISH) assay confirmed the existence of mono- and binuclear tetraploids (Fig. 2g, h). Overall, this indicated that both mononuclear and binuclear tetraploids were increased in siAnln vs. control livers and in control vs. siE2f8 livers.

**Figure 2.**
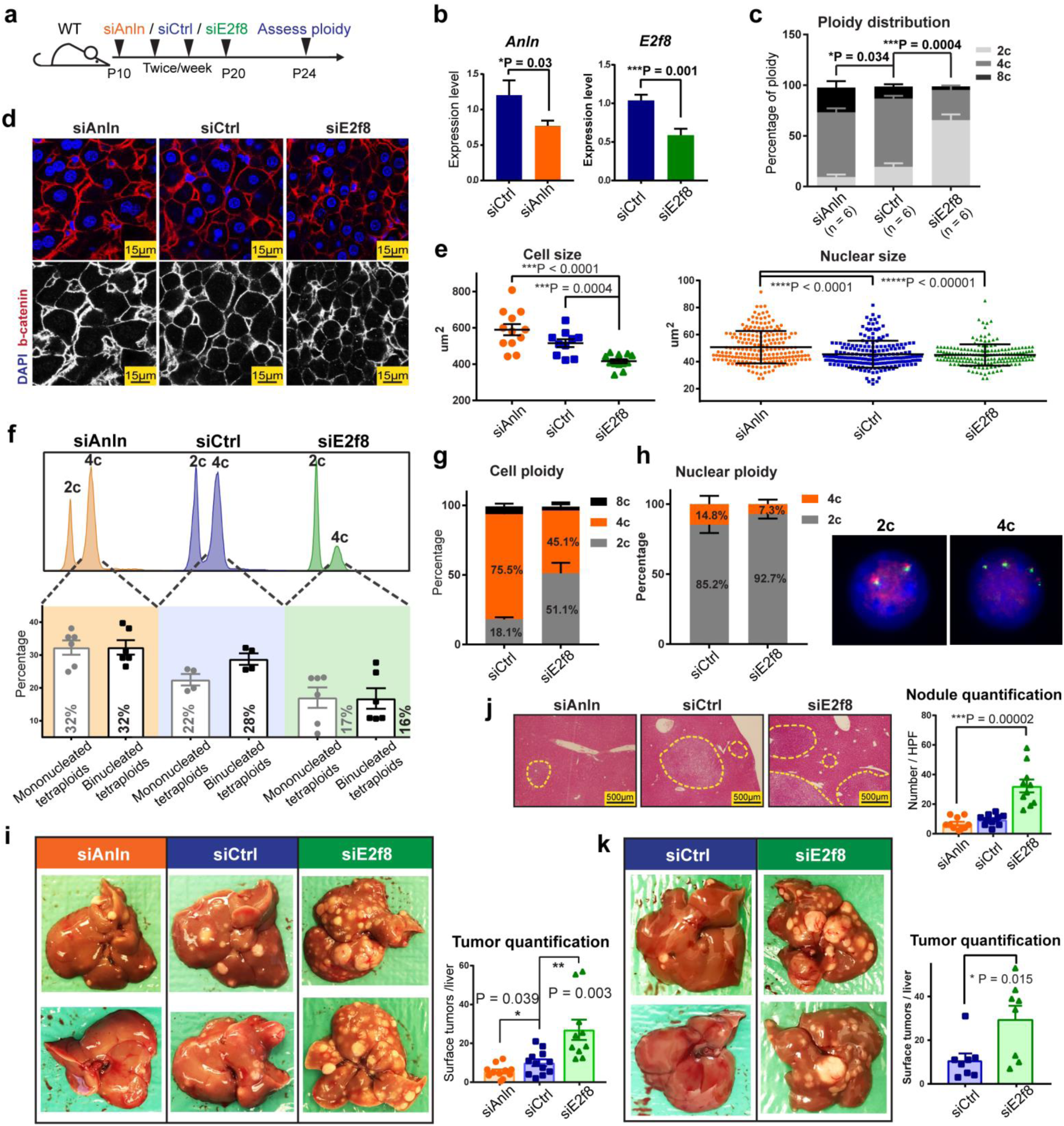
In vivo siRNAs that toggled ploidy revealed that the polyploid state protected against tumorigenesis. a. Schema for the siRNA experiment: *Anln*, *E2f8*, and scramble siRNAs (siCtrl) were encapsulated into lipid nanoparticles and injected into WT C3H strain mice starting at P10. Four total injections (two intraperitoneal and two retro-orbital) were performed twice per week. Between P24-26, hepatocytes were dissociated for ploidy analysis or mice were injected with DEN (75μg/g). Tumor burden was examined six months later.
b. mRNA knockdown after *Anln* and *E2f8* siRNA treatments. RT-qPCR was performed on the liver 4 days after the last siRNA dose.
c. Average cellular ploidy distribution within livers, as determined by PI staining and flow cytometry (n = 6 mice in each group were analyzed).
d. Confocal images of DAPI and Ctnnb1 stained siRNA treated livers. The bottom images highlight Ctnnb1 staining alone, marking membranes and cellular outlines.
e. Cross-sectional cell and nucleus size measurements. Each group includes 12 total analyzed image fields from 4 individual mice. For nucleus size quantification, each data point is one nucleus; for cell size quantification, each data point is an average of the cell sizes from one image and 3 images were taken from each mouse.
f. Representative cellular ploidy distribution of siRNA treated livers (upper panel). The lower panel shows the distribution of mono- and binucleated tetraploid hepatocytes. Calculations were performed as follows: % of mononucleated tetraploid cells = % tetraploid cells by FACS - % binucleated cells by confocal (images from Fig. 2D, n = 3 images were counted for each mouse, 2 mice for each group).
g. Cellular ploidy distribution of siRNA treated livers, as determined by Flow Cytometry (n = 2 mice/group).
h. On the left is the nuclear ploidy of the above siRNA treated livers, as determined by FISH. 60 nuclei were analyzed for each mouse. On the right are representative images showing a 2c and a 4c nucleus. 2 green and 2 red signals identify a 2c nucleus. 4 green and 4 red signals identify a 4c nucleus.
i. Gross tumor burden of siRNA treated livers harvested 6 months after DEN injection. Quantification of liver surface tumor numbers (right panel).
j. H&E liver histology showing tumor nodules circled by dashed lines (left panel). Quantification of microscopic nodules (right panel).
k. Tumor burden in mice treated with DEN at later time points after siRNA delivery. Mice were treated with four doses of siRNAs. 14 days after the last siRNA injection, one dose of DEN (100μg/g) was given, and 7.5 months later, tumor burden was assessed (n = 8 in each group). The liver surface tumors were quantified in the right panel.

Altered ploidy did not impact overall health, liver mass, body mass, hepatocyte differentiation, *CYP450* expression, or proliferation (Extended Data Fig. 2a-e). Moreover, we challenged these mice to acute regeneration assays such as partial hepatectomy and hepatotoxin treatments. Liver/body weight ratios after 2/3^rds^ partial hepatectomy were not significantly different between different ploidy groups (Extended Data Fig. 3a). No differences in necrosis and proliferation arose after one dose of DEN or carbon tetrachloride (CCl4) (Extended Data Fig. 3b, c). These results showed that polyploid and diploid hepatocytes were equivalently able to survive, recover, and proliferate after injuries. These findings are consistent with previous studies in mice with ploidy alterations^19,20^, again demonstrating that ploidy state has minimal influences on post-natal liver growth and regeneration, likely because only 2-4 cell division cycles are required for recovery after these profound, acute injuries.

Having established that inducible inhibition of *Anln* and *E2f8* allowed us to alter ploidy without introducing irreversible genetic mutations, we next evaluated the impact of ploidy on tumor development. We injected DEN (75μg/g x 1 dose) into mice four days after the last dose of siCtrl, siAnln, or siE2f8 (siRNA delivery schema in Fig. 2a). Six months later, siE2f8 treated hyperdiploid livers had significantly increased gross tumor burden, microscopic tumor nodules, and liver/body weight ratio than siCtrl livers (Fig. 2i, j and Extended Data Fig. 4a-c). In contrast, siAnln treated hyperpolyploid livers showed the opposite. Altogether, these findings suggested that the degree of liver polyploidy is inversely proportional to the efficiency of carcinogenesis. To further exclude the possibility of a residual siRNA effect, we also introduced DEN starting at 14, rather than 4 days, after the last siRNA injection. Again, polyploidy demonstrated a potent tumor suppressive effect (Fig. 2k and Extended Data Fig. 4d).

### *E2f8* knockout and inducible *Anln* shRNA mice protect against HCC in multiple models

Given the formal possibility that siRNAs or lipid nanoparticles could have off-target or nonspecific effects, we also used Cas9 to generate whole-body *E2f8* KO mice (Extended Data Fig. 5a), which have predominantly diploid hepatocytes in adulthood (Fig. 3a). Interestingly, we observed that *E2f8* WT, Het and KO livers had equivalent levels of ploidy at P15, and only diverged in ploidy state by P27 (Fig. 3a). We hypothesized that if ploidy is a specific and essential factor causing tumor suppression in *E2f8* KO livers, then mutagenizing and inducing cancer at P15 would not result in differences in HCC development, while inducing at P27 would result in large differences. Indeed, DEN given at P15 (25μg/g) resulted in no cancer differences between the three groups, while DEN at P27 (75μg/g) caused more HCCs in *E2f8* Het and *E2f8* KO mice when compared to *E2f8* WT mice (Fig. 3b-e, Extended Data Fig. 5b). These results further demonstrated that the *E2f8* gene itself contributed little to cancer development independent of the differences in ploidy seen at P27.

**Figure 3.**
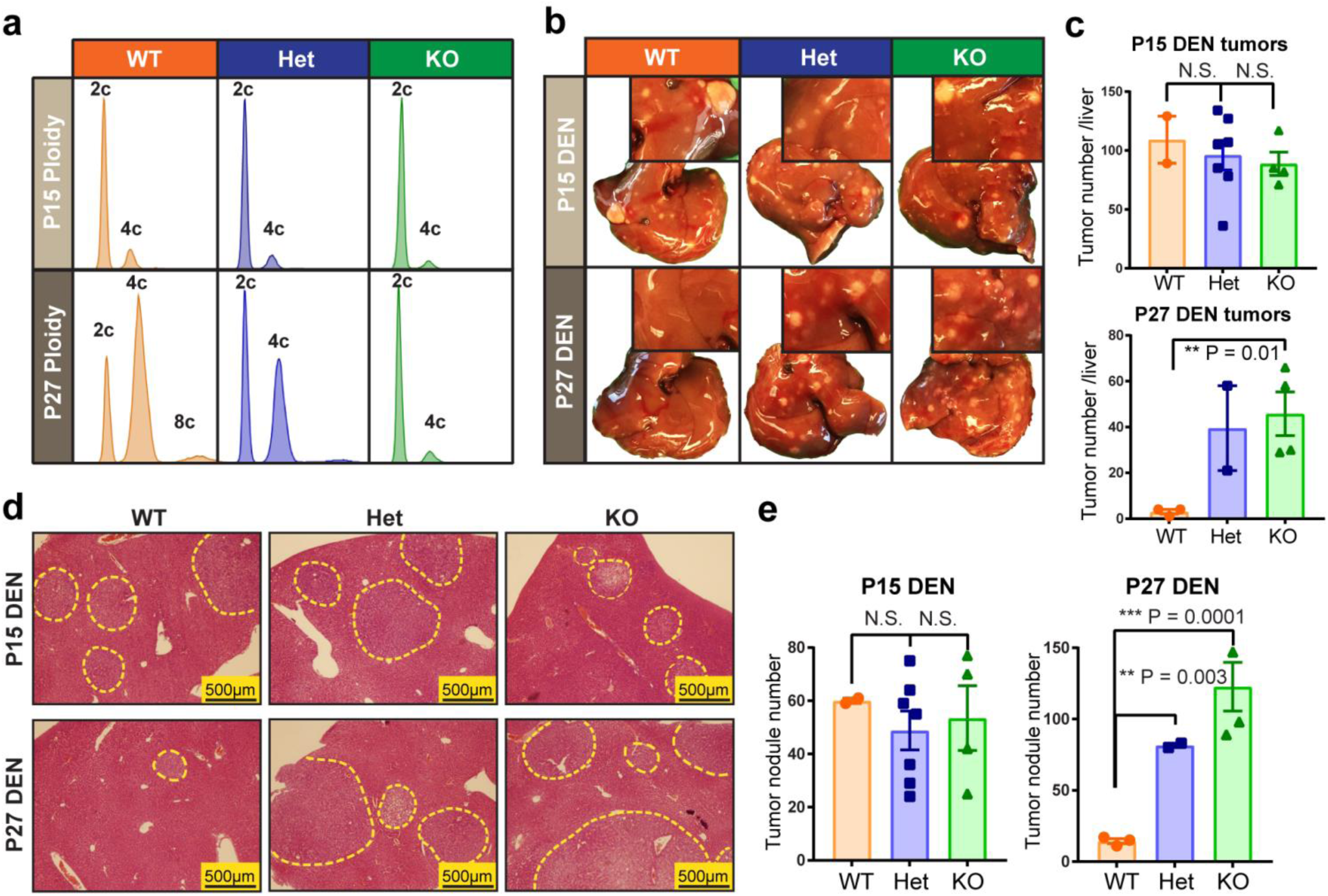
Reduced ploidy accounted for the increased tumorigenesis associated with *E2f8* deficiency. a. Representative cellular ploidy of *E2f8* WT, *E2f8* Het, and *E2f8* KO livers at P15 and P27, as analyzed by flow cytometry.
b. DEN was introduced by IP injection in one cohort at P15 (25μg/g) and in another cohort at P27 (75μg/g). Gross tumor burden was evaluated 4.5-5 months later. The inset images show higher magnification images that more clearly exhibit tumor burden.
c. Liver surface tumor quantification of *E2f8* WT, Het, and KO mice given DEN at P15 or P27.
d. Histology of *E2f8* WT, Het, and KO livers show microscopic tumor nodules (circled by dashed yellow outlines).
e. Quantification of tumor nodules per unit of cross sectional area.

Next, we wanted to generate a more potent and versatile mouse model to increase polyploidy. In addition, our goals were to drive greater levels of polyploidy and to employ a small inhibitory shRNA distinct from the siAnln used above in order to corroborate on target effects on *Anln*. Thus, we created a doxycycline (dox)-inducible transgenic mouse expressing an shRNA against *Anln*. Transgenic mice were derived from embryonic stem cells containing *Rosa-rtTA* and a GFP + sh*Anln* cassette under the control of a tetracycline responsive promoter element (TRE) (Extended Data Fig. 6a, b; transgenic design based on Scott Lowe’s group^26^). Dox could be used to induce *Anln* suppression in a temporally specific fashion (Fig. 4a). *Rosa-rtTa* alone or *Rosa-rtTa*; *TRE-shAnln* (hereafter called *Rosa* and *TG-shAnln*) transgenic mice exposed to dox water from P0-P20 showed normal growth, development, and liver function (Extended Data Fig. 6c-e). *Anln* mRNA levels were suppressed by 50% (Fig. 4b), which resulted in hyperpolyploid livers at multiple time points after dox withdrawal (Fig. 4c-e). These livers were similar to what was seen with *Anln* siRNA treatment, but had more profound levels of polyploidization. In addition, GFP protein (and likely *Anln* shRNA) completely disappeared by 15 days after dox withdrawal (Fig. 4f), demonstrating the reversibility of *Anln* suppression.

**Figure 4.**
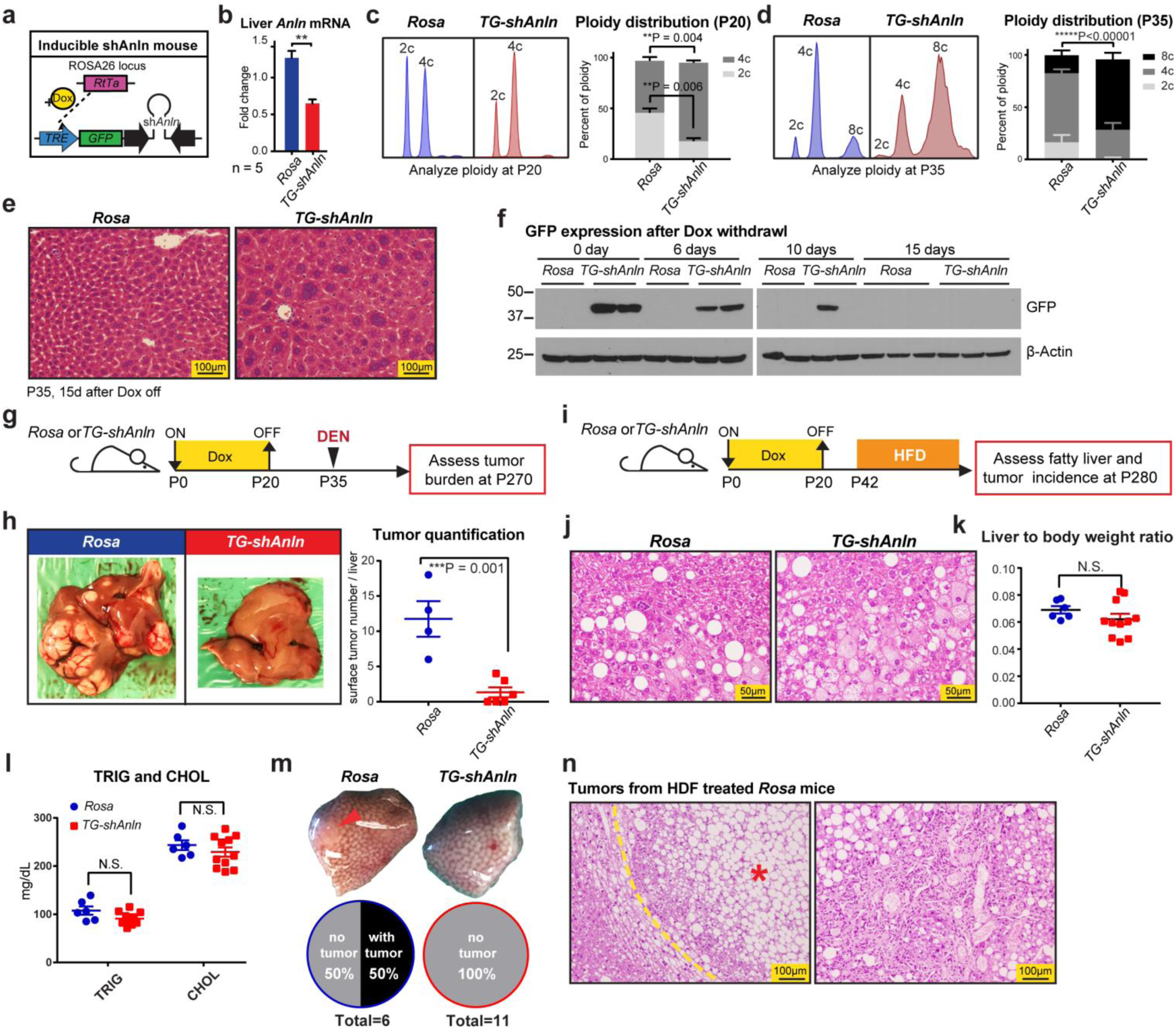
An inducible transgenic mouse model to temporally control polyploidization shows protection against DEN and high fat induced HCCs. a. An inducible double-transgenic mouse model carrying sh*Anln* cassette under the control of a tetracycline responsive promoter element (TRE). These mice carry a *Rosa-rtTA* knockin construct, allowing induction of *Anln* suppression with Doxycycline (Dox) water.
b. Transient Dox induction from P0 to P20 suppressed *Anln* mRNA levels in the liver.
c. Representative cellular ploidy distribution of *TG-shAnln* livers treated with dox from P0-P20 was determined by PI staining and flow cytometry at the age of P20 (left panel). On the right is the average ploidy distribution (n = 3 mice in each group).
d. Representative cellular ploidy distribution of *TG-shAnln* livers treated with dox from P0-P20 was determined by PI staining and flow cytometry at the age of P35 (left panel). On the right is the average ploidy distribution (n = 6 mice in each group).
e. H&E staining showing large, multinucleate hepatocytes in the induced group.
f. Western blots showing stability of GFP protein expression after dox induction and withdrawal.
g. Schema for the DEN induced HCC experiment in inducible shRNA mice: Transient dox treatment from P0-P20 in *Rosa-rtTa* or *TG-shAnln* mice established ploidy differences. At P35, mice were injected with DEN (100μg/g). Tumor burden was examined 8 months later.
h. Representative gross tumor burden from the DEN experiment. Quantitation on the right.
i. Schema for the high fat diet (HFD) induced HCC experiment in inducible *TG-shAnln* mice: Transient dox treatment mice established ploidy differences. Starting at P42, mice were given ad libitum HFD (60% calories from fat). Tumor burden was examined after 8 months.
j. H&E of *Rosa* and *TG-shAnln* HFD treated livers.
k. Liver to body weight ratios of HFD treated *Rosa* and *TG-shAnln* mice.
l. Triglyceride and cholesterol levels of *Rosa* and *TG-shAnln* mice after 8 months of HFD.
m. Representative gross tumor burden from the HFD experiment. 50% (3 of 6) of Rosa mice had tumors, while no (0 of 11) TG-shAnln mice carried tumor.
n. H&E of HFD induced tumors in *Rosa* control mice.

Hyperpolyploid mice derived from transient suppression of *Anln* were almost completely protected from DEN induced HCC development, confirming the siRNA results (Fig. 4g, h). We then wanted to exploit this model to analyze the function of polyploidy in a carcinogenesis model caused by another clinically important mechanism. Steatohepatitis represents an increasingly relevant risk factor for HCC and has been associated with an increase in polyploidy^8^. To induce long term fatty liver disease and HCC, we fed mice with high fat diet (HFD) after transiently inducing ploidy changes (Fig. 4i). After eight months, *Rosa* and *TG-shAnln* mice had similarly high levels of steatosis and liver function abnormalities (Fig. 4j-l and Extended Fig. 6f), but 50% of control mice while no hyperpolyploid mice developed HCC (Fig. 4m, n). In summary, multiple murine models increasing and decreasing ploidy corroborated the fact that higher levels of polyploidy suppressed HCC formation in diverse cancer models.

### Polyploids were protected from tumor suppressor LOH but not oncogene activation

To probe underlying mechanisms in the DEN HCC model, we first asked if ploidy significantly regulated metabolic properties that would influence tumorigenesis. DEN is first bioactivated by CYP450 family enzymes to become α-hydroxylnitrosamine^27^. Expression of CYP450 enzymes in general and zonation of Cyp2e1 in particular were unchanged in livers with distinct ploidy (Extended Data Fig. 3c and Extended Data Fig. 7a). Elevated levels of reactive oxygen species (ROS) secondary to hepatotoxins are known to accelerate tumor initiation^28^, but ROS levels were unchanged between siRNA treated mice and between *E2f8* WT/Het/KO mice, before and after DEN (Extended Data Fig. 7b, c). It is also known that following the DEN bioactivation, an ethyldiazonium ion is formed, binds DNA, and causes genotoxic damage^27.^ Though DNA damage markers such as p-Brca1, p-p53, and p-γH2A.X increased after DEN, the magnitude of induction was similar between groups (Extended Fig. 7d, e). Altogether, livers with altered ploidy did not exhibit differential xenobiotic metabolism, oxidative stress, or DNA damage responses.

Since the mutagenic activities of DEN were quantitatively similar, it was possible that polyploid cells were buffered from tumor suppressor loss but not oncogene activation. To isolate and test the impact of oncogene activation in mice with different levels of polyploidy, we overexpressed *MYC* using a liver specific driver (*LAP-tTa*) and a dox-inducible promoter (*TRE-MYC*) (Extended Data Fig. 8a)^29^. Prior to inducing *MYC* overexpression, we gave *LAP-tTA; TRE-MYC* mice four doses of siRNA to transiently alter ploidy (Fig. 5a). At P30, significant differences in ploidy, but not *MYC* expression levels, were observed (Extended Data Fig. 8b, c). Dox withdrawal at P25 leads to transformation of less than 1% of *MYC* expressing cells, a dynamic range that allowed us to sensitively quantitate the influence of ploidy on tumor initiation. Nine weeks post-induction, tumor development and liver to body weight ratios were indistinguishable between siRNA treated ploidy groups (Fig. 5b), showing that ploidy had little impact on *MYC* oncogene induced tumorigenesis.

**Figure 5.**
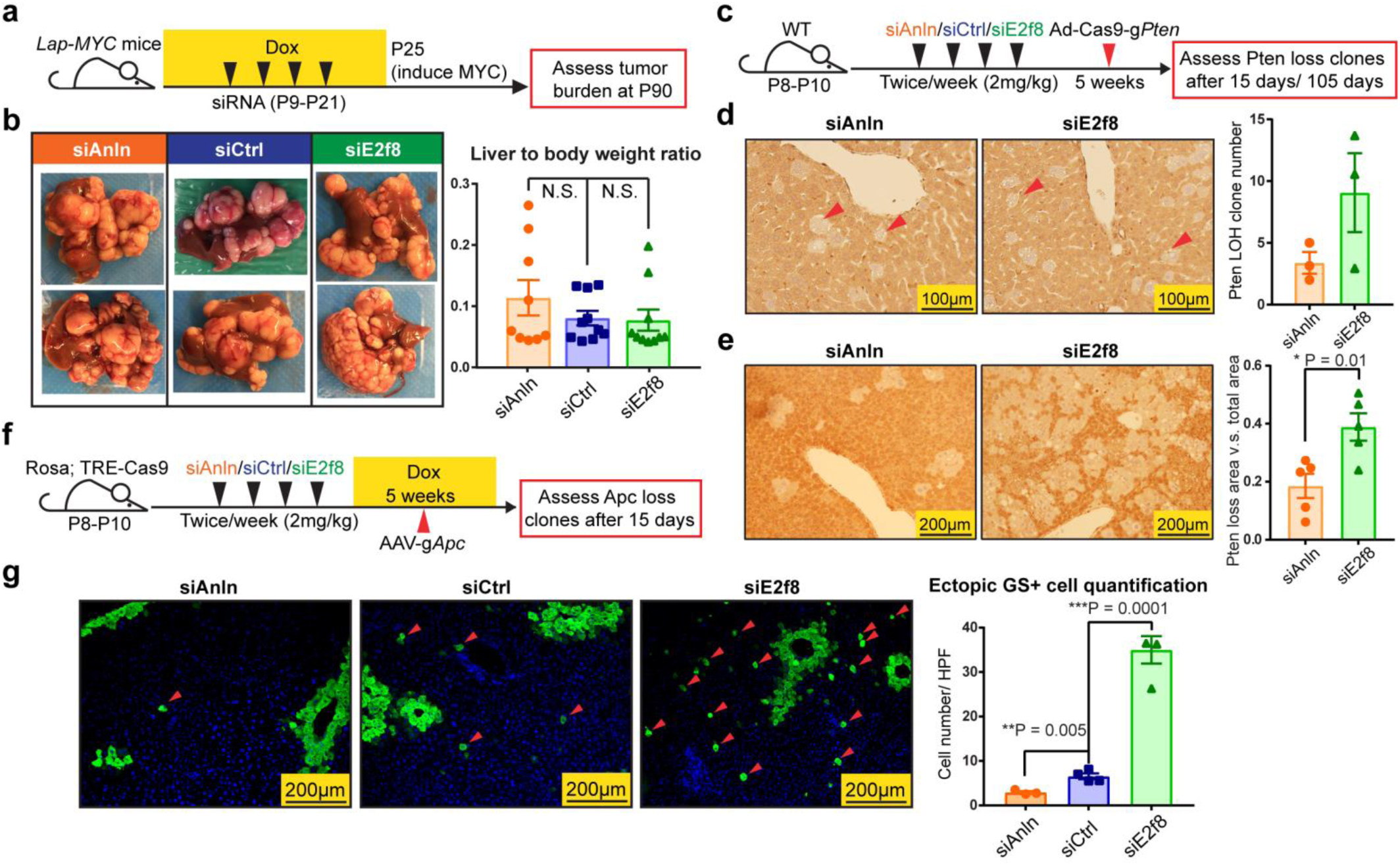
Polyploids were protected from tumor suppressor LOH but not oncogene activation. a. Schema of the *LAP-MYC* experiment: prior to inducing *MYC* overexpression by dox withdrawal at P25, *LAP-tTA; TRE-MYC* mice were given four doses of siRNA to transiently alter ploidy, as described in previous experiments. At P30, ploidy distribution and *MYC* expression levels were measured; At P90, tumor burden was assessed.
b. Tumor burden at P90 (left panel). Liver to body weight ratios, another metric of tumor burden, are shown in the right panel.
c. Schema of the Ad-*Cas9-sgPten* experiment: WT mice were given four doses of siRNA to alter ploidy; two weeks later, adenovirus carrying Cas9 and a guide strand RNA targeting *Pten* (Ad-*Cas9-sgPten*) was injected into these mice (10^9^ pfu/mouse); 15 days or 110 days later, LOH was assessed by Pten immunohistochemistry.
d. Pten staining (left panel) on siAnln and siE2f8 treated livers at 15 days after virus injection. Red arrowheads point to the cells with *Pten* deletion, and LOH is quantified in the right panel.
e. *Pten* immunohistochemistry (left panel) in siAnln and siE2f8 treated livers 110 days after Adenovirus injection. *Pten* deleted area vs. total cross sectional area was quantified (right).
f. Schema of the AAV-sg*Apc* experiment: Dox inducible *Rosa; TRE-Cas9* mice were given four doses of siRNA to alter ploidy. Five days later, dox (1g/L) was given to induce Cas9 expression. Two weeks after the last dose of siRNA, AAV-*sgApc* (5 x 10^12^ pfu/mouse) was retro-orbitally delivered; 15 days later, the *Apc* LOH were assessed by Glutamine Synthetase (GS) immunofluorescence.
g. GS staining (left panel) 15 days after AAV-sg*Apc* injection. Red arrowheads point to ectopic GS positive cells that result from homozygous *Apc* deletion. Note the normal, non-ectopic GS staining that surrounds central veins (CV) that was not quantified. Ectopic GS positive cells per imaging area were quantified in the right panel.

It remained possible that the impact of DEN-induced oncogene activation was agnostic to ploidy, but that DEN induced tumor suppressor loss of heterozygosity (LOH) is more difficult to achieve in hepatocytes with wholesale genome duplications. To quantitate the dynamics of tumor suppressor loss, we engineered an adenovirus carrying Cas9 and a guide strand RNA targeting *Pten* (Ad-*Cas9-sgPten*) (sgRNA was validated in reference^30^), a commonly inactivated tumor suppressor gene in HCC. 15 days after IV Ad-*Cas9-sgPten* delivery into mice with different levels of ploidy, *Pten* LOH was assessed with immunohistochemistry (Fig. 5c). The frequency of hepatocytes with complete *Pten* deletion was inversely proportional to the extent of polyploidy (Fig. 5d, Extended Data Fig. 8d). After 110 days, this effect was even more pronounced (Fig. 5e, Extended Data Fig. 8e). To confirm that the degree of ploidy did not change susceptibility to adenoviral infection, we verified that equal numbers of cells expressed GFP after Adenovirus-GFP delivery (Extended Data Fig. 8f). To rule out the possibility of a *Pten* specific-phenomenon, we also assessed *Apc*, a tumor suppressor in the WNT pathway. We injected an Adenoassociated virus carrying a guide strand against exon 8 of *Apc* (AAV-*sgApc*) into dox-inducible *Cas9* mice (*Rosa-rtTa; TRE-Cas9*) subjected to siRNA induced ploidy changes (Fig. 5f). In vitro, this particular *sgApc* effectively mutagenized *Apc* (Extended Data Fig. 8g) and had previously been used to generate *Apc* null hepatocytes^31^. To quantitate the number of *Apc* null clones in vivo, we probed for ectopic Glutamine Synthetase (GS), a specific and sensitive marker of aberrant WNT activation in the liver^32^. Strikingly, hyperdiploid livers were much more susceptible to *Apc* LOH than control livers, and hyperpolyploid livers harbored the fewest GS+ cells (Fig. 5g). These data support the concept that polyploid livers are protected from tumor suppressor loss and are not more sensitive to oncogene activation.

Reduced susceptibility to tumor suppressor LOH paired with reduced tumorigenesis in hyperpolyploid livers suggested that DEN-induced HCCs were in large part dependent on tumor suppressor loss. Previously, DEN was shown to preferentially select for oncogenic mutations (*Hras* and *Ctnnb1*) in studies focused on identifying mutations in these pathways^33,34^. We aimed to more broadly and thoroughly map the mutational landscape of these tumors, so we sequenced 242 of the most commonly mutated genes in human and murine HCC in 50 individual DEN induced tumors (Fig. 6a, Extended Data Table 2). We did identify a core group of recurrent, mutually exclusive mutations in oncogenes such as *Egfr* (Phe254Ile), *Hras* (Gln61Arg), and *Braf* (Val637Glu), but a majority of the most commonly mutated genes were bona fide tumor suppressors such as *Mll2* (*Kmt2d*), *Brca2*, *Arid1a*, *Atm*, *Apc*, and *Tsc2*. Overall, these data suggest that DEN tumor transformation depends on tumor suppressor loss in addition to EGFRRAS-MAPK pathway activation, supporting the idea that tumor protection in polyploids is in part mediated through retention of WT tumor suppressor alleles.

**Figure 6.**
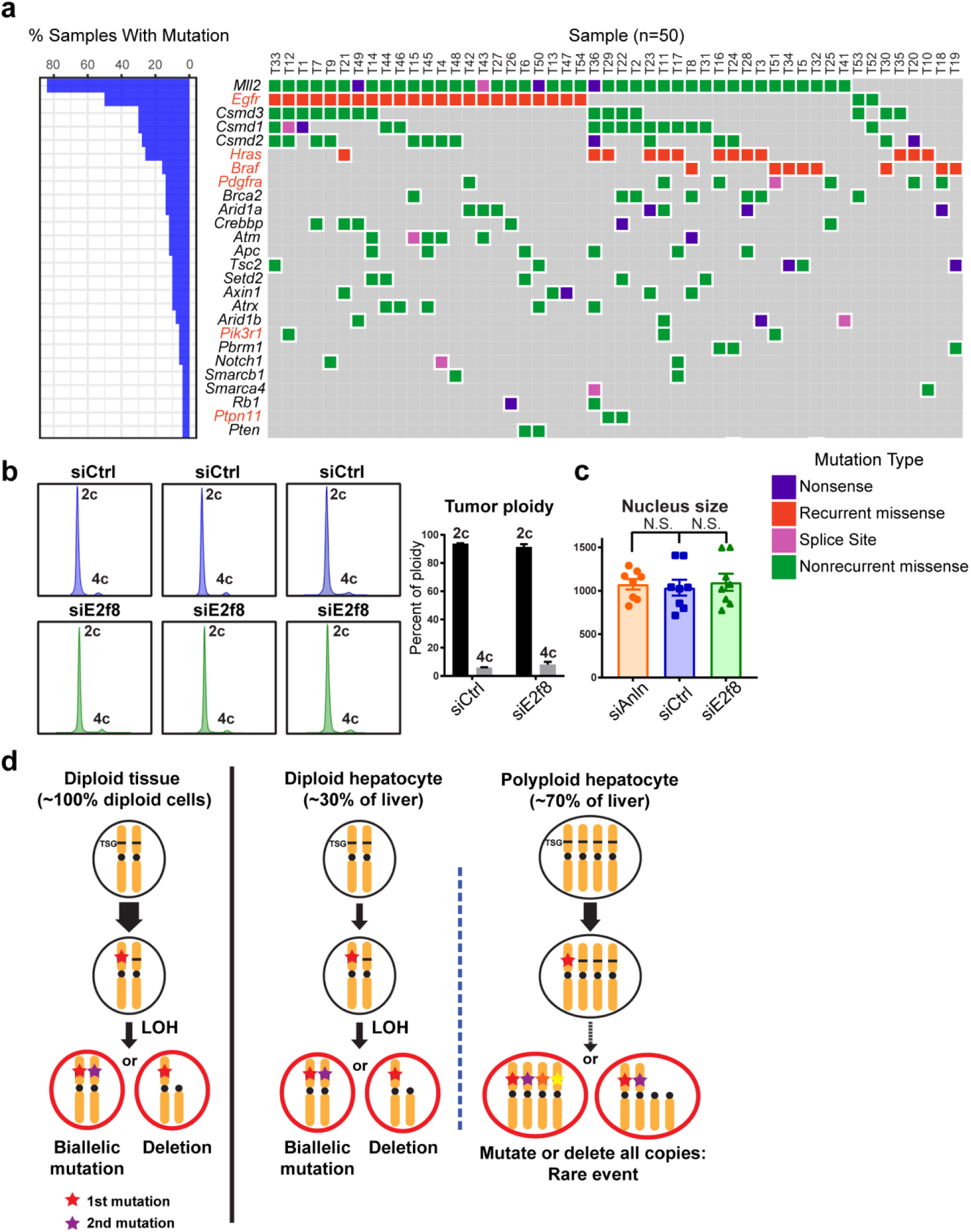
Tumor suppressor mutations are prevalent in HCCs from diploid and polyploid mice. a. Targeted sequencing on 50 DEN induced HCC tumors. 242 of the most commonly mutated genes in human and murine HCC were sequenced. Genes highlighted in red are oncogenes and those in black are tumor suppressors.
b. Cellular ploidy distribution of DEN induced HCC tumors from siRNA treated mice. Data is averaged on the right (tumor n = 10).
c. Nucleus sizes of DEN induced HCC tumors from siRNA treated mice. The nucleus size was quantified using Photoshop based on H&E staining images. 4 mice in each group were quantified, 2 tumors per mouse.
d. This model depicts how tumor suppressors could be lost in diploid and polyploid cells. Note that the liver contains both diploid and polyploid cells.

Given that the mammalian liver is comprised of a mixture of polyploid and diploid cells, our work would predict that HCCs more likely originate from diploid cells. If this was true, it would follow that HCCs are also more frequently diploid rather than polyploid. We found that even tumors arising from siCtrl mice with polyploid cells were predominantly diploid (Fig. 6b, c). In human HCCs examined by flow cytometry, 80% of HCCs were comprised of diploid cells^35^^-^^37^. Altogether, this suggests that human HCCs more likely arise from diploid cells, consistent with the idea that the polyploid state is less compatible with cancer development.

## Discussion

The fact that some animals and organs are polyploid, with cells containing whole genome duplications, has perplexed scientists for decades. It is known that 50-80% mammalian hepatocytes are polyploid, and conditions such as chronic hepatitis, steatohepatitis, and oxidative stress have all been associated with increased polyploidy^8,38^, but it is unknown if ploidy changes represent compensation, cause, or bystander effect of such disease states. Analysis of genetically engineered knockout mice with hyperpolyploid or hyperdiploid livers have not been able to unequivocally link ploidy with cellular fitness, regeneration, or cancer^18,20^. In part, this is because the reagents used to assess the response of ploidy states to injuries usually confound the effects of ploidy with the effects of the prominent cell cycle genes used to change ploidy.

Here, we devised multiple in vivo methods to transiently alter ploidy in a reversible fashion, such that long-term consequences of such changes could be assessed and compared. In agreement with previous studies^11,20^, we found that polyploidy had little impact on acute injury or regeneration, but chronically, the polyploid state demonstrated tumor suppressive functions in multiple cancer models. It is interesting that the extent of ploidy increase or decrease in different siRNA, GEMM, and inducible transgenic models appeared to correlate linearly with tumor number and burden. Moreover, we believe that this protection is due in large part to buffering against tumor suppressor loss rather than limiting proliferation after oncogenic insults (Model shown in Fig. 6d). This is consistent with human data indicating that most HCCs are diploid^35^^-^^37^.

We believed that it was essential to use mouse models to interrogate the role of ploidy in cancer because it is the most rigorous way to examine tumor initiation. For example, in vivo and in vitro human HCCs, which are already transformed, cannot be used to study ploidy’s role in the transition state between normal and malignant cells. In addition, we used cancer models that were the most reflective of human disease states. The DEN model generates cancers that are not dependent on singular, artificially strong genetic drivers such as *Yap* or *Akt*. Using expression profiling, Snorri Thorgeirsson showed that DEN induced mouse HCCs were more similar to poor survival human HCCs than other mouse cancer models^39^. Also, Allan Balmain’s group recently argued for the use of chemically induced mouse cancer models because they more accurately reflect the diverse genomics of human cancers^40^. Indeed, the tumors in our study harbored a wide diversity of oncogenic and tumor suppressor mutations that reflect the biology of human HCCs. The DEN model may mechanistically and genetically reflect the type of liver cancers that arise in humans exposed to mutagens such as aflatoxin, which is a frequent HCC-inducing contaminant of foods in tropical and sub-tropical climates in Africa, Asia and South America^41^. It is conceivable that this unique feature of the liver in part evolved to mitigate the cancer risk associated with these types of common liver mutagens, which frequently induce lethal cancers during the reproductive years. In addition, we showed that polyploid livers are also resistant to high fat induced HCC. These data suggest that the polyploidization associated with non-alcoholic steatohepatitis may represent a protective and tumor suppressive stress response.

In future studies, it would be interesting to determine if there are specific diploid populations in the liver that have greater cancer risk and if this risk can be modulated with ploidy manipulations. Previously, Wang et al. identified a “stem-like” hepatocyte population around the central vein^14^. This population of cells repopulated the liver in the absence of injury and were 60% enriched for diploid cells, but it is unknown if the diploid or polyploid cells within this Axin2 compartment have greater self-renewing capacity or tumorigenic potential. It is possible that increasing ploidy in this population might change regenerative capacity or cancer risk. Given that most hepatocytes are capable of extensive proliferation^11,42^, it is also possible that other diploid populations, whether or not they are stem cells, could respond to distinct liver injuries and contribute to cancer formation. In this study, we aimed to answer a fundamental question about polyploidy in liver biology, but it is possible that transient or persistent polyploidzation not only serves physiologic functions but could be exploited therapeutically.

## Methods

### Mice

All mice were handled in accordance with the guidelines of the Institutional Animal Care and Use Committee at UTSW. *E2F8* KO mice were made by the Children’s Research Institute Mouse Genome Engineering Core. Cas9 was used to target intron 1 and 3, which led to the deletion of exons 2 and 3. (see Table S1 for sgRNA sequences). These mice were made and maintained in the C3H/HeJ background. *TG-shAnln* embryonic stem cells were injected by the UTSW Transgenic Core. The strategy was based on work from Scott Lowe’s group ^26^. Briefly, the *shAnln* sequences were cloned into the pCol-TGM vector (Addgene #32715) and electroporated into KH2 embryonic stem cells together with a Flpe vector. Validated KH2 clones were then injected into blastocysts to generate knockin transgenic mice. All other mice used in experiments were wild-type C3H/HeJ mice. All experiments were done in an age and sex controlled fashion unless otherwise noted in the figure legends. The experimental mice were randomized using simple randomization method.

### Cell Culture and Transfection

H2.35 (ATCC) immortalized hepatocytes were cultured in DMEM supplemented with 4% (vol/vol) FBS, 1x Pen/Strep solution (Thermo Scientific HyClone) and 200nM Dexamethasone (Sigma). Cells were transfected with 25pmol siRNA (Life Technologies) in 6-well plates by using Lipofectamine RNAiMAX and OptiMEM (Life Technologies), as described in the manufacturer’s instructions. Transfected cells were cultured for 72 hours before performing RNA extraction and fixation for flow cytometry. The cell line used in this study has been tested as mycoplasma free.

### Partial hepatectomy

Surgery was performed as previously described ^43^.

### High fat diet experiments

High fat diet (HFD) was purchased from RESEARCH DIETS INC. 60% calories were from fat. Animals were kept on HFD from 1.5 to 8 months of age.

### Chemical injury experiments

CCl4 is diluted 1:10 in corn oil (Sigma), and administered IP at a dose of 0.5 ml/kg of mouse as described previously ^44^. Diethylnitrosomine (DEN, Sigma) is diluted in saline and administered IP at different doses, depending on mouse age. For P19 mice, 25μg/g of mouse was injected once; for P24-P27 mice, 75μg/g was injected once; for P34 mice, 100μg/g was injected once.

### Virus experiments

100μL of Ad-*Cas9-sgPten* (Vectorbiolabs) was retro-orbitally injected at a dose of 10^9^ pfu/mouse. 100μL of Adenovirus-GFP (University of Iowa Viral Vector Core) was retro-orbitally injected at a dose of 1.4 X 10^8^ pfu/mouse. 100μL of AAV-sgApc (5 X 10^12^ pfu/mouse) was retro-orbitally injected.

### RNA Extraction and RT-qPCR

Total RNA was isolated using Trizol reagent (Invitrogen). For qRT-PCR, cDNA synthesis was performed with 1μg of total RNA using iScript Reverse Transcription Kit (Biorad). See Table S1 for primers used in these experiments. Gene expression levels were measured using the ΔΔCt method as described previously ^45^.

### Western Blot Assay

Mouse liver tissues were homogenized and lysed in T-PER Tissue Protein Extraction Reagent (Thermo Scientific Pierce). Western blots were performed in the standard fashion. The following antibodies were used: anti-β-Actin (Cell Signaling, #4970), Anti-Anln (Abcam 154337), Anti-E2f8 (Abcam 109596), Anti-p-BRCA1 (Cell Signaling, #9009), Anti-p-p53 (Cell Signaling, #9286), Anti-rabbit IgG, HRP-linked Antibody (Cell Signaling, #7074) and Anti-mouse IgG, HRP-linked Antibody (Cell Signaling, #7076)

### Histology, Immunohistochemistry, and Immunofluorescence

Tissue samples were fixed in 4% paraformaldehyde (PFA) and embedded in paraffin. In some cases, frozen sections were made. Immunohistochemistry was performed as previously described ^45^. Primary antibodies used: Ki67 (Abcam, ab15580), Cyp2e1 (Abcam, ab28146), Pten (Cell Signaling, #9559S), Ctnnb1 (BD Transduction Laboratories #610154), p-γH2A.X (Cell Signaling, 2577S). For immunohistochemistry, detection was performed with the Elite ABC Kit and DAB Substrate (Vector Laboratories), followed by Hematoxylin Solution counterstaining (Sigma). For immunofluorescence, second antibody Alexa Fluor 594 goat anti-mouse IgG1 (life technologies) was used. Confocal images were taken by Zeiss LSM 780 Upright confocal/multiphoton microscope. H&E slides were interpreted by a clinical pathologist with expertise in human liver cancer diagnosis.

### Primary hepatocytes isolation

Primary hepatocytes were isolated by two-step collagenase perfusion. Liver perfusion medium (Thermo Fisher Scientific, 17701038), liver digest medium (Thermo Fisher Scientific, 17703034) and Hepatocyte wash medium (Thermo Fisher Scientific, 17704024) were used. Cell number and viability were determined by Trypan blue exclusion in a hemocytometer.

### Flow Cytometry

For detection of ploidy populations, primary hepatocytes (2x10^6^/mL) or transfected H2.35 cells (1 ×10^6^ mL) were fixed in 75% ethanol at -20°C, then incubated with 500μL (2x10^6^ mL) of PI/Rnase Staining Buffer (BD Pharmingen) at 25°C for 15 minutes. Cells were analyzed with a BD FACS Aria Fusion machine (BD Biosciences). For Ki67 staining, cells were fixed in 75% ethanol at -20°C, washed twice in staining buffer (PBS with 1% FBS), and incubated with anti-Ki67 antibody (BD#550609, 1:50) at room temperature (RT) for 30 minutes in the dark. Then the cells were incubated with secondary antibody Alexa Fluor 488 goat anti-mouse IgG1 (life technologies) for 20min at RT, and resuspended in PI/Rnase Staining Buffer (BD Pharmingen), and then flow cytometry analyzed.

### Fluorescence In Situ Hybridization (FISH) assay

Two FISH probes for mChr12 were provided by Dr. Hongtao Yu lab. Bacterial artificial chromosomes were nick translated with the CGH Nick Translation Kit (Abbott Molecular) using Red 580 dUTP and Green 496 dUTP (Enzo Life Sciences). Isolated primary hepatocytes were treated with 0.56% KCL and fixed with cold 3:1 methonal: acetic acid solution. Then the fixed cells were dropped onto slides, and allowed to dry. Then the slides were incubated with probes, covered with coverslips sealed with rubber cement. Slides were incubated at 80°C for 5 min and 37°C overnight. Following incubation, slides were washed in 0.5XSSC+0.1% at 70°C for 5 min, 1XSSC for 3X5 min at room temparature, 4XSSC+0.1% Tween for 5 min at room temperature, and 2XSSC for 5 min, and then mounted with DAPI (Vectorlabs). Samples were analyzed and scored under a Olympus IX83 microscope. For analysis of nuclei ploidy, 2 green and 2 red dots was counted as a 2c nucleus, and 4 green and 4 red dots was counted as a 4c nucleus. 60 nuclei were counted for each sample.

### Genomic DNA isolation and sequence processing

50 flash-frozen tumors free of visible normal tissue were used for library preparation. The genomic DNA was extracted using QIAGEN AllPrep DNA/RNA Kit (Cat. 80204). Integrity of genomic DNA was assessed by electrophoresis on 1% agarose gels, and concentration was determined by nanodrop. gDNA was sonicated into 500bp fragments and purified using Genomic DNA Clean & Concentrator Kit (ZYMO RESEARCH). The DNA library was prepared using Ovation Target Enrichment System (NuGEN) following the manufacturer’s instructions. Target genes were selected from human HCC and mouse sequencing studies ^46^^-^^48^ and some well-known cancer related genes. The target probes were synthesized by NuGEN. More than 99.5% of the probes had >90% coverage. The sequencing was performed using a 150 bp single-end protocol on Illumina NextSeq 500 platform.

### Sequence processing

BCL files from Illumina Nextseq 500 sequencing were converted to FASTQ files by bcl2fastq (Illumina). After trimming by trim_galore package, BWA-MEM (version 0.7.15) was used to align FASTQ files to reference genome GRCm38 with subsequent processing by Samtools (version 1.3) and Nudup.py (Nugen) to ensure proper file formatting and remove duplicates. Alignments were then recalibrated and realigned by GATK (version 3.5). We acquired 42 million uniquely mapped reads on average for the 53 samples we sequenced with an average on target coverage at 128X and more than 86.6% region has more than 50X coverage.

### Identification and annotation of somatic SNVs

To detect somatic variants in tumor samples, we use the somatic variant detection program Mutect (version 1.1.7). 50 tumor samples were called against a panel of 3 normal liver samples from two WT C3HHeJ mice to the reference genome GRCm38. Variants passed the Mutect high-confidence somatic mutation filters were selected and further filtered against from known SNVs in C3HHeJ mice provided by http://ftp-mouse.sanger.ac.uk (mgp.v3). SNVs were further annotated using snpEff (version 4.2) and Variant Effect Predictor. SNVs predicted to have High or Moderate impact were compared to a stringent list of 125 driver genes in human cancer by Vogelstein et al ^49^ to map putative driver genes and mutations. The mutation landscape was graphed by using GenVisR (a package from R Bioconductor) waterfall plot algorithm on putative driver genes for all 50 tumor samples (Fig. 6a).

### Statistical analysis

The sample size was determined by producing a confidence interval estimate with a specified margin of error to ensure that a test of hypothesis has a high probability of detecting a meaningful difference in the parameter. The data in most figure panels reflect multiple experiments performed on different days using mice derived from different litters. Variation is indicated using standard error presented as mean ± SEM. The variances of two groups that are being compared are similar. Two-tailed Student’s *t*-tests (two-sample equal variance) were used to test the significance of differences between two groups. Statistical significance is displayed as *p* < 0.05 (*) or *p* < 0.01 (**) unless specified otherwise. The investigators were blinded during the processes of mice treatments and data analysis. Image analysis for the quantification of cell proliferation, *Pten* deletion, GFP+ cells, liver surface tumor and malignant nodule numbers were performed in a blinded fashion.

### Data availability

The DEN tumor DNA targeted sequencing data was reported in Figure 6a and Extended Data Table 2. The source dataset for the data reported in this Figure will be made available at publication or earlier if requested.

## Acknowledgments

We thank H. Yu, M. Buszczak, S. Morrison, and H. Sadek for critical input and advice; P. Gopal for pathological interpretation; E. Choi for assistance with FISH; Jian Xu and the CRI sequencing core for genome sequencing. H.Z. was supported by the Pollack Foundation, an American Cancer Society pilot grant, a NIH/NCI R01 grant (1R01CA190525), a Burroughs Welcome Career Medical Award, and a CPRIT New Investigator grant.

## Author contributions

S.Z. and H.Z. conceived the project, performed the experiments and wrote the manuscript. L.L., S.Z., L.N., assisted with experiments and mouse husbandry. K.Z. and D.S. assisted with the chemistry experiments. Y.Z. helped to make the mouse model. X.L. performed bioinformatics analysis.

